# The novel SCN5A-P1891A mutation is associated with left ventricular hypertrabeculation and links Na_v_1.5 to cardiomyocyte proliferation and disrupted 3D cardiac tissue formation

**DOI:** 10.1101/2025.02.19.639199

**Authors:** Qasim A. Majid, Chandra Prajapati, Xiaonan Liu, Julia Kari-Koskinen, Lotta Pohjolainen, S. Tuuli Karhu, Heikki Ruskoaho, Markku Varjosalo, Katriina Aalto-Setälä, Virpi Talman

**Author notes:** These authors contributed equally to this work (Qasim A. Majid, Chandra Prajapati). <u>Correspondence:</u> Virpi Talman Faculty of Pharmacy P.O. Box 56 (Viikinkaari 5E) FI-00014 University of Helsinki Finland E.

## Abstract

**Background:** Left ventricular hypertrabeculation (LVHT) is a heterogenous cardiac condition with a complex and poorly understood aetiology. We comprehensively characterised the effect of a novel P1891A mutation in the *SCN5A* gene, which encodes the voltage-gated sodium channel Na_v_1.5, identified in a Finnish family diagnosed with LVHT.

**Methods:** We generated *SCN5A-*P1891A mutation-carrying human induced pluripotent stem cell-derived cardiomyocytes (P1891A-hiPSC-CMs) and performed electrophysiological assessments, including patch-clamp studies, and fluorescent calcium imaging, to determine the mutation’s effect on hiPSC-CM electrophysiology. We also evaluated the impact of the mutation on the proliferative capacity in response to mitogenic stimuli and on the hypertrophic response following cyclic mechanical stretch or endothelin-1 treatment. Further, we assessed the effect on contractile parameters in three-dimensional (3D) contractile hydrogels (engineered heart tissues, EHTs) and conducted advanced proteomics to understand the consequences of the mutation on Na_v_1.5 protein-protein interactions.

**Results:** The *SCN5A-*P1891A mutation reduced the sodium current densities and increased both the sodium window current and arrhythmogenicity; however, action potential parameters were unaffected. Advanced proteomics characterised, for the first time, the complete Na_v_1.5 interactome and revealed that the *SCN5A-*P1891A mutation negated interactions with fibroblast growth factor 12 (FGF12) and FGF13, that are known to modulate sodium channel activity. Baseline proliferation was unchanged, although aged P1891A-hiPSC-CMs demonstrated enhanced proliferative capacity following mitogenic stimulation. Further, P1891A-hiPSC-CMs exhibited a heightened stress response upon mechanical stretch, resulting in the upregulation of heart failure-associated genes. Strikingly, EHTs derived from P1891A-hiPSC-CMs yielded disparate phenotypes. Whilst the majority condensed only partially and failed to beat synchronously, a small subset condensed fully yet exhibited weak contractile properties, alongside age-associated functional decline. In contrast, EHTs derived from healthy control hiPSC-CMs consistently condensed fully and demonstrated a positive correlation between post-fabrication age and contractile properties

**Conclusions:** Our study presents a unique aetiology of LVHT and reveals a novel association between *SCN5A* mutations and enhanced human cardiomyocyte proliferation. Further, the inability of P1891A-hiPSC-CMs to consistently form fully condensed 3D cardiac tissues may be linked to their abnormal response to mechanical stretch and provides a powerful 3D model for future mechanistic research and drug development studies to better understand and treat LVHT.

## Introduction

Left ventricular hypertrabeculation (LVHT), traditionally referred to as left ventricular noncompaction cardiomyopathy (LVNC), is a poorly understood cardiac disorder characterised by a thin compact myocardial wall and excessive trabeculae, which are interspersed by deep intertrabecular recesses and extend into the LV cavity ^1^. LVHT is often diagnosed by examining the ratio of trabecular to compact myocardium, with a ratio exceeding 2.0–2.3 typically considered indicative of the condition ^2^. The clinical presentation of LVHT is highly heterogeneous, ranging from asymptomatic patients to those presenting with arrhythmia, thromboembolic events, and LV dysfunction. Further, LVHT can manifest as either an isolated phenotype or in conjunction with other cardiac diseases ^3^. The ramifications of confounding factors, as well as the correlation between the degree of excessive trabeculation and subsequent major adverse cardiac events, such as sudden cardiac death and heart failure, are equally complex. Indeed, initial disease presentation and a thinner compact layer appear to be more detrimental to longer term prognosis than the extent of hypertrabeculation ^4^, which, in asymptomatic individuals, is not correlated with left ventricular ejection fraction (LVEF) ^5^.

Hypertrabeculation may result from an imbalance in the proliferative capacity of the compact and trabecular layers of the developing human myocardium, disrupting their ratio and potentially explaining the varied effects of cardiomyocyte (CM) proliferation reported in LVHT development ^6, 7^. However, the precise molecular mechanisms underpinning this are not fully understood, and LVHT may also present as a reversible phenotype in response to environmental factors that lead to excessive preload, such as during pregnancy ^8^ or intense exercise ^9^.

LVHT is largely considered an inherited disease ^10^, often arising from autosomal dominant transmission ^2^, although X-linked recessive mutations have also been documented ^11^. At least 189 genes have been reported to be associated with LVHT ^12^. Among these, genes involved in protein degradation (*MIB1*) ^7^ and sarcomere function (*MYH7*) ^13^ have been shown to have causative effects whilst mutations in genes encoding for the ion channels *HCN4* ^14^ and *RYR*2 ^15^ have also been implicated in LVHT.

The *SCN5A* gene encodes for the pore-forming alpha subunit of the voltage-gated sodium channel, Na_v_1.5, that generates the cardiac sodium current (I_Na_) and is thus responsible for the rapid upstroke of the cardiac action potential (AP) ^16^. Owing to its complex structure, comprised of four structurally homologous domains that form the channel pore, the location of mutations within this structure vastly influences their effect on the channels function and gating properties. In turn, *SCN5A* mutations have been associated with a plethora of cardiac diseases, including Long QT syndrome Type 3 (LQT3), arising as a result of gain-of-function (GOF) mutations, and Brudgada syndrome (BrS) resulting from loss-of-function mutations (LOF) ^17^. *SCN5A* mutations have also been attributed to the arrhythmogenic pathophysiology of LVHT, with affected patients experiencing a greater number of arrhythmias and a higher rate of heart failure than LVHT patients who do not have *SCN5A* mutations ^18^. However, to date, no direct causal relationship between *SCN5A* mutations and LVHT has been established.

Recently, we have identified a Finnish family with LVHT. Genetic screening revealed a single, novel P1891A missense mutation within *SCN5A* in family members exhibiting the LVHT phenotype. To ascertain the effect of this mutation on Na_v_1.5 electrophysiology and CM functionality in 2D and 3D models, we generated human induced pluripotent stem cells (hiPSCs) from the most clinically affected family member and differentiated these into hiPSC-CMs (P1891A-hiPSC-CMs). Further, as Na_v_1.5 functionality is heavily influenced by the proteins with which it interacts ^19^, we used the recently developed multiple approaches combined (MAC)-tagged proteomics workflow ^20^ to investigate the protein interactome of both wild type (WT) Na_v_1.5 and Na_v_1.5 p.P1891A, revealing novel interacting proteins and the mutation-driven loss of known interactors. Together, this extensive characterisation of P1891A-hiPSC-CMs and Na_v_1.5 proteomics identifies, for the first time, a potential causative effect of *SCN5A* mutations in LVHT, further highlighting the significance of this ion channel beyond generating the sodium current.

## Methods

### Data availability

The data explaining the major findings of our study are shared in the main article and in the Supplemental Material. Please refer to the Supplemental Materials and Methods for detailed experimental methods. Research materials utilised in this study are outlined in the Supplemental Materials and Methods and are included in the Major Resources Table in the Supplemental Material. The mass spectrometry dataset generated in this study is available from the MassIVE database (https://massive.ucsd.edu/) with web access MSV000095720. Specific data sets generated during the study are available from the corresponding author upon reasonable request.

### Clinical information

The proband, a 42-year-old Finnish female (patient III-1, Figure 1A) demonstrated hypertrabeculation of the left ventricle with normal LVEF, as analysed with echocardiography and cardiac MRI. Mild hypertrophy was also observed, with septal thickness of 13-15 mm, without hypertension or valvular disease. Palpitations were occasionally reported, and monomorphic ventricular extrasystoles (VES) were observed following a 24-hour Holter recording, covering a maximum of 8% of beats over 24 hours and approximately 2% per 24 hours when receiving the beta blocker, bisoprolol. Non-sustained ventricular tachycardias (NSVT) were also observed prior to beta blocker medication. On genetic screening, the proband was observed to carry a novel, single nucleotide missense mutation in the *SCN5A* gene, located on chromosome 3, causing a proline to alanine substitution at position 1891 *(p.P1891A c.5568C>G)* in the C-terminal intracellular region of the protein (Figure 1B).

**Figure 1.**
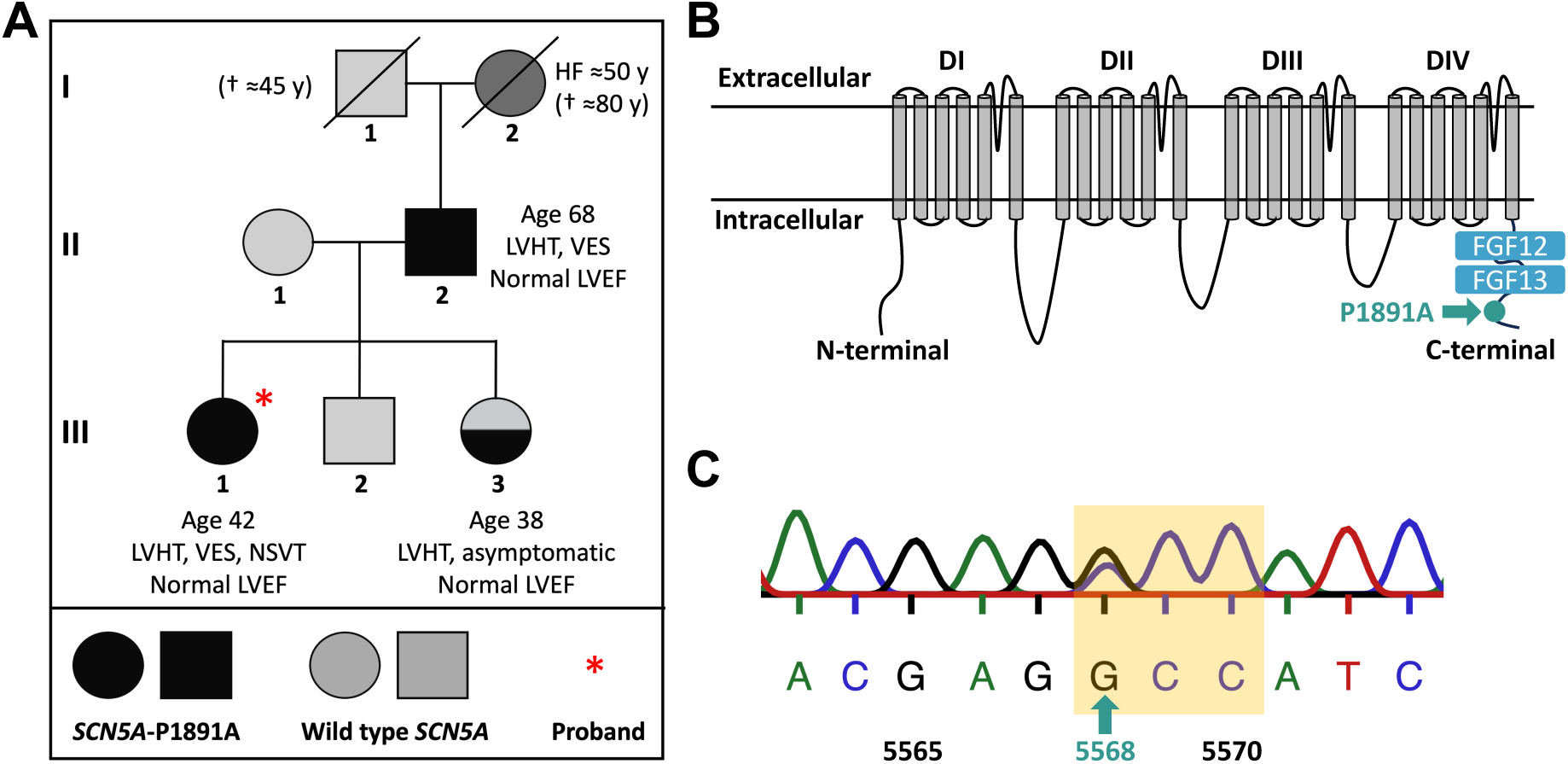
Clinical presentation of LVHT in a Finnish family. (A) Pedigree of the Finnish family. Males and females are denoted by squares and circles, respectively. Black symbols depict patients diagnosed with LVHT. The diagonal line represents a deceased family member, and the proband is highlighted with a red star. (B) Schematic diagram illustrating the β-subunit of Na_v_1.5. The binding sites for FGF12 and FGF13 are highlighted alongside the location of the P1891A mutation (green arrow). (C) Sanger sequencing confirming the *SCN5A c.5568C>G* mutation (green arrow) within the mutation-carrying hiPSCs that changes the codon (yellow box) and results in the Na_v_1.5 p.P1891A mutation.

In the probands family history, unexplained heart failure (patient I-2) had been observed at the age of 50 years (Figure 1A). The cardiac structure and function of the proband’s siblings and parents were also analysed. The probands female sibling (III-3) and father (II-2) also had excessive trabeculation, whist her mother (II-1) and brother (III-2) exhibited normal cardiac structure. Despite the LVHT diagnosis, the sister was asymptomatic; however, the father had similar palpitations and VES as the proband. On genetic screening, both the father and the sister were determined to carry the same *SCN5A*-P1891A mutation as the proband, whereas her mother and brother did not.

## Results

### Differentiation of P1891A-hiPSC lines generated P1891A-hiPSC-CMs that displayed enhanced arrhythmogenicity and reduced sodium current density

To characterise the effect of this novel *SCN5A*-P1891A mutation on cardiomyocyte function, we first reprogrammed the dermal fibroblasts isolated from the proband and generated two P1891A-hiPSC clones: UTA.14301.*SCN5A*, and UTA.14303.*SCN5A*. The emerging hiPSCs retained the *SCN5A*-P1891A mutation, as determined by Sanger sequencing (Figure 1C), and expressed markers associated with pluripotency (Figure S1A-B) as well as genes indicative of the three germ layers (Figure S1B), but not those associated with the hiPSC reprogramming factors (Figure S1C). Further, the two hiPSC lines exhibited normal karyotypes (Figure S1D) and short tandem repeat (STR) profiles in line with the dermal fibroblasts (Figure S1E-F).

Following spontaneous differentiation of both P1891A-hiPSC clones, we performed perforated patch-clamp on spontaneously beating hiPSC-CMs to examine the cardiac AP. The P1891A-hiPSC-CMs demonstrated enhanced arrhythmogenicity relative to healthy control hiPSC-CMs (Figure 2A-C), consistent with the proband’s clinical presentation (patient III-1, Figure 1A). Assessment of intracellular calcium (Ca^2+^) dynamics through Ca^2+^ imaging further substantiated this increased arrhythmogenicity (Figure 2D-F).

**Figure 2.**
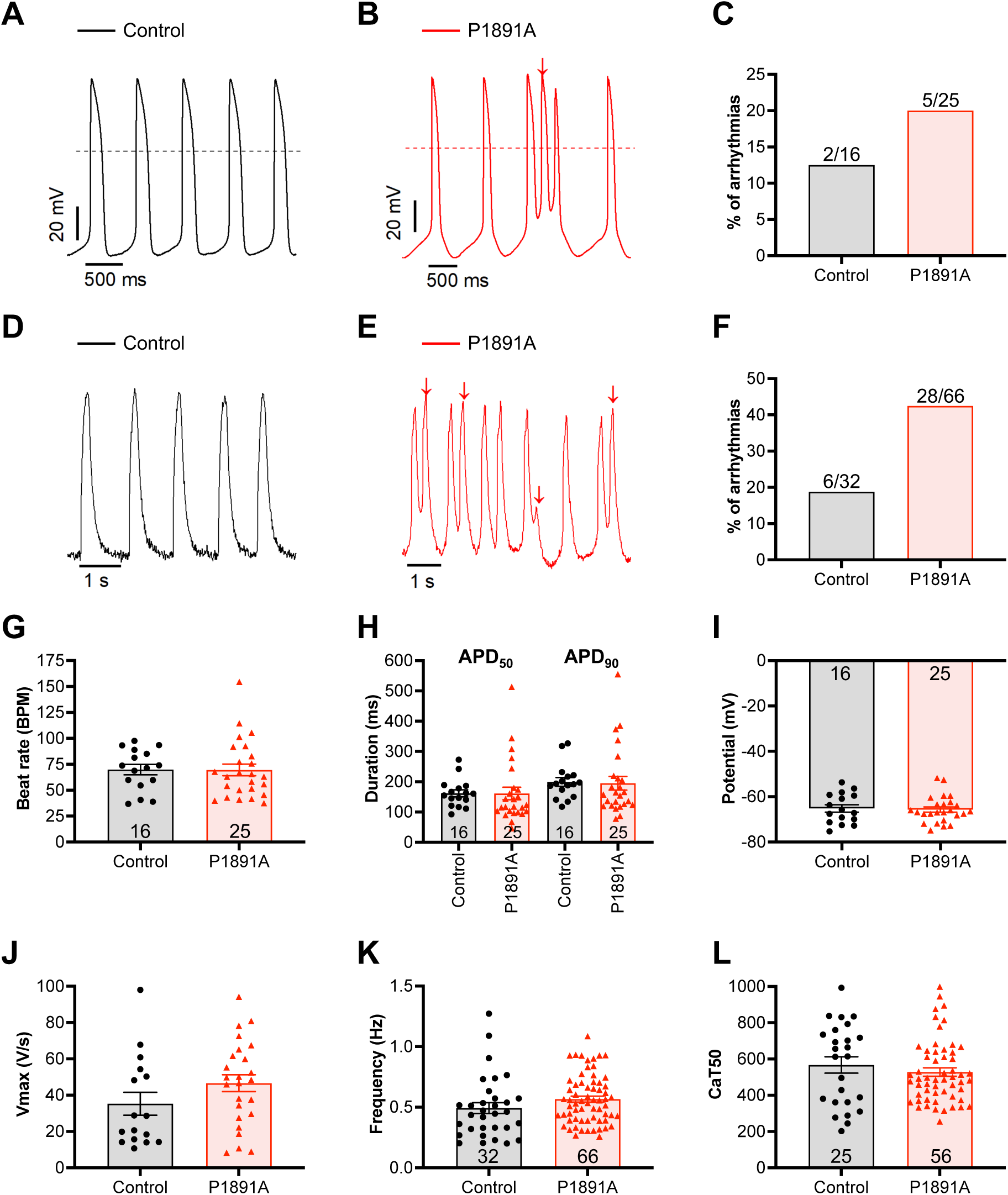
P1891A-hiPSC-CMs recapture the arrhythmogenic nature of the proband and exhibited a reduced sodium current density. Representative action potential recordings from **(A)** healthy control- and **(B)** P1891A-hiPSC-CMs. The red arrow indicates arrhythmia. **(C)** The percentage (%) of arrhythmogenic events detected from healthy control- and P1891A-hiPSC-CMs via the patch-clamp technique. Representative traces of intracellular calcium dynamics from **(D)** heathy control- and **(E)** P1891A-hiPSC-CMs. Arrhythmias are highlighted by the red arrows. **(F)** The percentage (%) of arrhythmogenic events detected within healthy control- and P1891A-hiPSC-CMs via calcium imaging. Quantification of the **(G)** beat rate, **(H)** action potential duration (APD) at 50% and 90% of repolarisation, **(I)** maximum diastolic potential, and **(J)** the maximum upstroke velocity of the action potential of healthy control- and P1891A-hiPSC-CMs. Comparison of the **(K)** frequency and **(L)** mid-calcium transient duration of healthy control- and P1891A-hiPSC-CMs as determined via intracellular calcium imaging. **(C and F)** The values above the bars represent the fraction of hiPSC-CMs that exhibited arrhythmogenic events. **(All)** The N numbers for each experiment are displayed within the respective graph with N referring to the number of cells analysed. Data are presented as mean ± SEM.

However, additional electrophysiological parameters determined from spontaneous AP recordings including beats per minute (BPM) (Figure 2G), AP durations at 50% and 90% repolarisation (APD_50_ and APD_90_, respectively) (Figure 2H), maximum diastolic potential (MDP) (Figure 2I), and maximum upstroke velocity (Vmax) (Figure 2J) were unchanged in the P1891A-hiPSC-CMs relative to healthy control hiPSC-CMs. Similarly, we found that the frequency of the intracellular Ca^2+^ transients (Figure 2K) and the mid-Ca^2+^ transient duration (CaT50, Figure 2L), as ascertained via Ca^2+^ imaging, were comparable to healthy control hiPSC-CMs.

To further investigate the electrophysiological parameters, we injected current through the patch pipette and stimulated the hiPSC-CMs at 1 Hz to achieve a consistent beating rate. The APD at -20mV and -60mV, as well as the Vmax, were unchanged between healthy control- and P1891A-hiPSC-CMs (Figure S2).

Next, we conducted whole-cell patch-clamp studies to investigate the effect of this *SCN5A*-P1891A mutation on the sodium (Na^+^) channel function. Upon assessment of the sodium current (I_Na_) traces (Figure 3A), we noted that the P1891A-hiPSC-CMs exhibited significantly decreased I_Na_ densities between –40 to –20 mV (Figure 3B). The time-dependent I_Na_ inactivation kinetic, as represented by τ_fast_ and τ_slow_, were similar between healthy control and P1891A-hiPSC-CMs (Figure 3C). In addition, the half-maximal activation potential (V_1/2_) and the slope factor of voltage dependence of (in)activation (k) from both I_Na_ activation and inactivation did not differ statistically between control and P1891A-hiPSC-CMs (Figure 3D, SI Table 1). Notably, a higher fraction of Na^+^ channels activated inappropriately in P1891A-hiPSC-CMs between the test potential of around -70mV to -30mV, as observed in the inactivation curve (Figure 3D). This increased the Na^+^ window current, defined as the area beneath the point at which the two curves intersect, in P1891A-hiPSC-CMs (Figure 3D, inset). However, similar time constant values (SI Table 2) suggested no difference in the ability of the channel to recover from I_Na_ activation (Figure 3E) nor the kinetics of entry into the slow inactivation state (Figure 3F).

**Figure 3.**
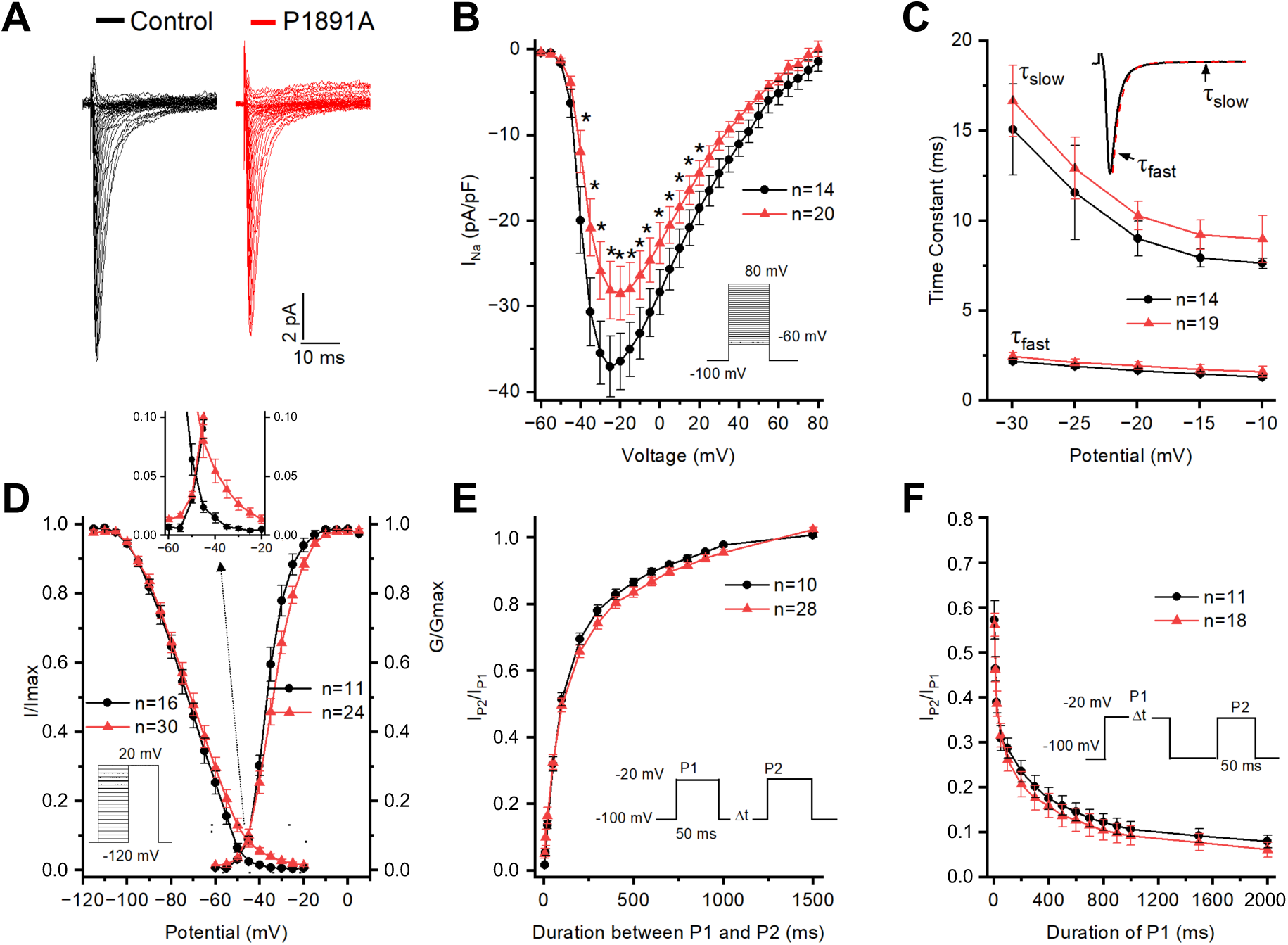
P1891A-hiPSC-CMs exhibit a reduced sodium current density. **(A**) Representative sodium current (I_Na_) traces from healthy control-(black trace) and P1891A-hiPSC-CMs (red trace). **(B)** Quantification of the normalised current density-voltage relationship of I_Na_. The inset shows the voltage clamp protocol. Data were analysed with an unpaired T-test: *P<0.05. **(C)** Averaged fast (τfast) and slow (τslow) time constants of I_Na_ inactivation plotted as a function of membrane potential. Inset: the trace marked in black was fitted with a two-phase exponential decay function shown in red. **(D)** The voltage dependence of steady-state activation and inactivation was plotted as a function of membrane potential and fitted to single Boltzmann functions. Inset: The voltage dependence of steady-state activation and inactivation between –60 mV and –20 mV from which the Na^+^ window current was determined. **(E)** Time-course of recovery after inactivation. Peak I_Na_ elicited by P2 was normalised (P2/P1) and plotted as function of the recovery interval. **(F)** The time-course of entry into the slow inactivation state. Peak I_Na_ elicited by P2 was normalised (P2/P1) and plotted as function of the duration of P1. **(E-F)** The inset shows the voltage clamp protocol. The N numbers for each experiment are displayed within the respective graph with each N referring to a single hiPSC-CM. Data are presented as mean ± SEM.

As the *SCN5A*-P1891A mutation occurs upstream of a calmodulin binding site that modulates the gating properties of the Na^+^ channel ^21^, we next investigated the calcium current (I_Ca_) via whole-cell patch-clamp. There were no differences in the I_Ca_ density (Figure S3A) nor the voltage dependence of activation (Figure S3B) suggesting these parameters were not impinged upon by the *SCN5A*-P1891A mutation. We also investigated the size of the hiPSC-CMs via cell capacitance and similarly found no statistically significant difference between healthy control and P1891A-hiPSC-CMs (Figure S3C).

Thus, whilst the P1891A-hiPSC-CMs recaptured the arrhythmogenic nature of the proband, we determined the wider electrophysiological effect of this mutation to be relatively minor. Therefore, we conducted further characterisation extending beyond electrophysiological parameters to determine the effects of this mutation and help contextualise its role in LVHT aetiology.

### *SCN5A* expression was upregulated in P1891A-hiPSC-CMs

We next sought to investigate whether the P1891A-hiPSC-CMs differed from control hiPSC-CMs in terms of their intracellular localisation of Na_v_1.5, and basal expression levels of *SCN5A* mRNA and cardiac stress-responsive genes. We employed a well-established small-molecule directed differentiation protocol ^22^ to attain highly pure (> 95%) populations of healthy control- and P1891A-hiPSC-CMs as visualised by immunofluorescent staining for the cardiomyocyte marker, α-actinin (Figure 4A and 4B). As in healthy control hiPSC-CMs, Na_v_1.5 colocalised with α-actinin in the P1891A-hiPSC-CMs (Figure 4B), with no significant difference in the average intensity of Na_v_1.5 staining in either the nucleus or perinuclear area (Figure S4).

**Figure 4.**
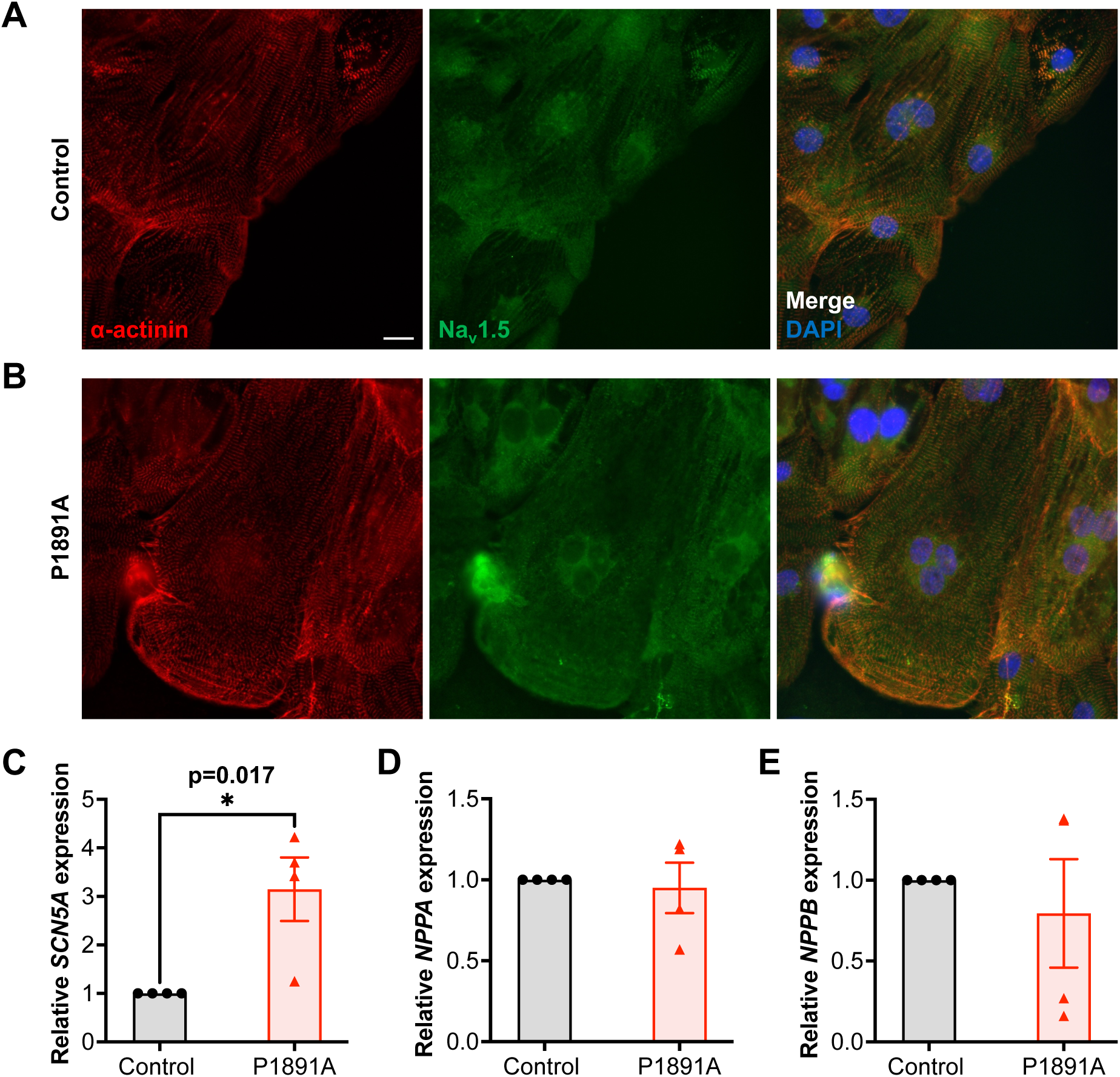
*SCN5A* expression is upregulated in P1891A-hiPSC-CMs. Representative immunofluorescence images of **(A)** healthy control- and **(B)** P1891A-hiPSC-CMs stained for the cardiomyocyte marker α-actinin (red), Na_v_1.5 sodium channel (green), and the nuclear marker DAPI (blue). Scale bar represents 25 µm. qPCR analysis of **(C)** *SCN5A*, **(D)** *NPPA*, **(E)** and *NPPB* gene expression in healthy control-(black bar) and P1891A-hiPSC-CMs (red bar). Data are normalised to healthy control hiPSC-CMs and presented as mean ± SEM, N=4 independent hiPSC-CM differentiations. Data were analysed with an unpaired T-test: * P<0.05.

We also determined baseline gene expression of *SCN5A* mRNA, which was significantly upregulated in the P1891A-hiPSC-CMs (Figure 4C). The basal expression of *NPPA* and *NPPB*, which encode for the stress-responsive natriuretic peptide A (ANP) and natriuretic peptide B (BNP), respectively, was unchanged relative to healthy control hiPSC-CMs (Figure 4D and 4E), indicating that the P1891A-hiPSC-CMs did not exhibit signs of cardiac stress under unstimulated conditions.

### The *SCN5A*-P1891A mutation perturbed Na_v_1.5-FGF protein-protein interactions

Na_v_1.5 protein-protein interactions (PPIs) are essential to the function of the Na^+^ channel ^23^, and are partly mediated by its subcellular localisation, further highlighting the complexity of this protein, which has been shown to invoke biological effects beyond its roles in generating the I_Na_ and the cardiac AP ^24^, ^25^. Therefore, to comprehensively determine PPIs at both the N- and C-terminals of the WT Na_v_1.5 and how these are affected by the *SCN5A*-P1891A mutation, we performed a recently established Multiple Approaches Combined (MAC)-tag workflow ^20, 26^. By combining affinity purification with mass spectrometry (AP-MS) and proximity-dependent biotinylation identification (BioID) (Figure 5A), we identified 43 high-confidence interactions (HCIs) that were consistently detected by both techniques (Figure 5B, 5C, and SI Table 3). These novel Na_v_1.5-interacting proteins were overwhelmingly involved in biological processes occurring within either the mitochondria, the endoplasmic reticulum (ER), or the Golgi apparatus (including TMEM87A).

**Figure 5.**
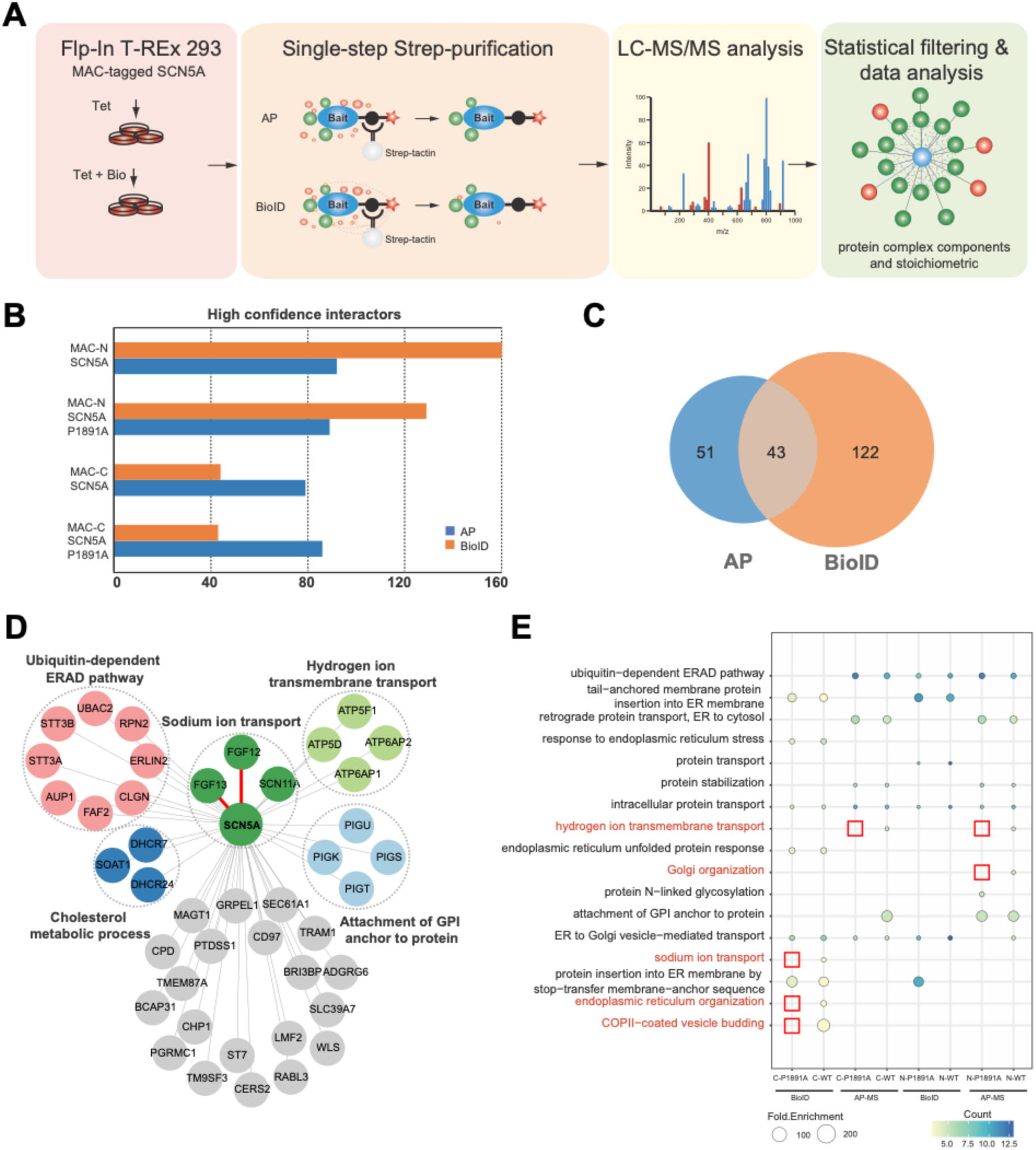
Na_v_1.5 protein-protein interactions with FGF12 and FGF13 are disrupted by the *SCN5A*-P1891A mutation. (A) Schematic overview of the MAC-tag workflow employed to determine the physical and functional connections formed by Na_v_1.5. Isogenic, inducible Flp-In-T-REx 293 cell lines are generated via transfection with either N-terminal or C-terminal MAC-tagged wild type (WT) *SCN5A* or *SCN5A*-P1891A. (B) The number of high-confidence interactions (HCIs) identified for WT *SCN5A* and *SCN5A*-P1891A at the N- and C-terminals via AP-MS (blue bars) and BioID-MS (orange bars). (C) Venn diagram illustrating the HCIs identified disparately by the two approaches as well as the 43 HCIs identified by both approaches that were used to construct (D) the protein interaction network of Na_v_1.5. The red lines denote protein-protein interactions that are disrupted by the *SCN5A-*P1891A mutation. The coloured nodes represent the different biological functions the interacting proteins are associated with. (E) Gene Ontology (GO) term enrichment analysis of the 43 identified HCIs. The scale denotes the percentage of interactors expressed per GO term (the count) and the fold enrichment (the size of the dot). GO terms that are not identified for the *SCN5A*-P1891A mutation relative to WT *SCN5A* are highlighted in red.

Intriguingly, we identified fewer HCIs for the Na_v_1.5 p.P1891A mutation relative to WT Na_v_1.5 (Figure 5B). This suggested that, in addition to disrupting specific interactions, including those with fibroblast growth factors 12 (FGF12) and 13 (FGF13) (Figure 5D), which are known to modulate the inactivation parameters of the Na^+^ channel ^27, 28^, the P1891A mutation broadly diminished the overall interactome of Na_v_1.5 (Figure 5B). This reduced capacity of Na_v_1.5 p.P1891A to form stable and functional protein complexes may impinge upon the non-electrophysiological roles of the Na^+^ channel.

To establish these processes, we conducted gene ontology (GO) term enrichment analysis, which revealed that the WT Na_v_1.5 HCIs were involved in processes including hydrogen ion transmembrane transport, Golgi organisation, Na^+^ ion transport, endoplasmic reticulum organisation, and budding of COPII-coated vesicles, which were not predicted for Na_v_1.5 p.P1891A (Figure 5E).

### P1891A-hiPSC-CMs exhibited enhanced proliferative capacity upon stimulation with mitogenic compounds

Owing to the contrasting data on the proliferation of CMs within the trabecular and compact layers of the LVHT myocardium ^6, 7^, we next assessed the proliferative capacity of P1891A-hiPSC-CMs relative to healthy control hiPSC-CMs up to 6-weeks post-differentiation through a BrdU incorporation assay (Figure 6A and 6B). Under baseline conditions, we found the proliferative capacity of P1891A-hiPSC-CMs to be comparable to that of healthy control-hiPSC-CMs at 3, 4, and 5 weeks post-differentiation (Figure 6C). Whilst we noted a two-fold increase in the percentage of proliferating P1891A-hiPSC-CMs as compared to healthy control-hiPSC-CMs at 6 weeks post-differentiation, this was not statistically significant (Figure 6C). Subsequently, we treated the two hiPSC-CM lines with a combination of the GSK-3β inhibitor CHIR99021 and the p38 MAPK inhibitor SB203580, both of which have been reported to induce CM proliferation ^29, 30^. Although these treatment regimens did not enhance the proliferation of young (less than 5-weeks post-differentiation) P1891A-hiPSC-CMs, the combinatorial treatment strategy invoked marked proliferation of 6-week-old P1891A-hiPSC-CMs, contrasting with age-matched healthy control hiPSC-CMs (Figure 6D). This suggested that the *SCN5A*-P1891A mutation may influence the proliferation of CMs in LVHT, albeit in a more nuanced manner requiring inhibition of certain pathways than previously investigated disease mediators that exert their effect in utero ^6, 7^.

**Figure 6.**
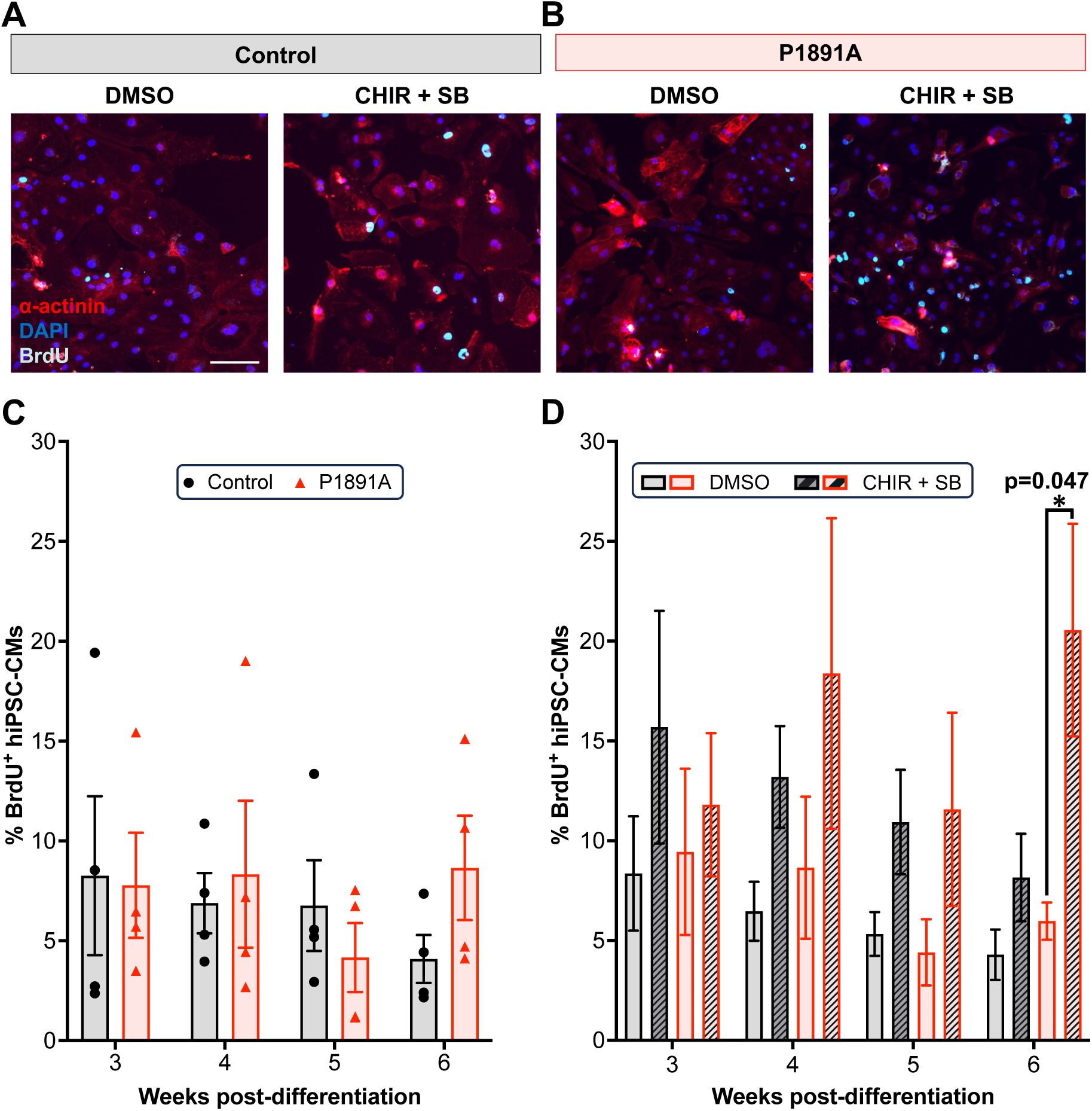
P1891A-hiPSC-CMs demonstrate enhanced proliferation upon stimulation with mitogenic compounds. Representative immunofluorescence images of 6-week post-differentiation **(A)** healthy control- and **(B)** P1891A-hiPSC-CMs cultured in RB^+^ medium containing BrdU and either 0.1% DMSO (vehicle control) or 5 μM CHIR99021 (CHIR) and 1 μM SB203580 (SB) for 24 hours. hiPSC-CMs were stained for the cardiomyocyte marker α-actinin (red), the proliferation marker BrdU (cyan), and the nuclear marker DAPI (blue). Scale bar represents 100 µm. The percentage (%) of BrdU^+^ healthy control-(black-bordered bars) and P1891A-hiPSC-CMs (red bars) under **(C)** basal conditions and **(D)** following treatment at 3-, 4-, 5-, and 6-weeks post-differentiation. Data are presented as mean ± SEM, N=3-4 independent hiPSC-CM differentiations. The data was analysed with a two-way ANOVA followed by a Bonferroni’s multiple comparison test: * P<0.05.

### P1891A-hiPSC-CMs demonstrated an increased stress-response upon mechanical but not endothelin-1 induced stress

The mechanosensitivity of Na_v_1.5 allows for an upregulation in I_Na_ and an enhancement in channel kinetics in response to mechanical stretch ^31^. Further, given that LVHT can be reversibly acquired under excessive preload ^8, 9^, we subjected healthy control- and P1891A-hiPSC-CMs to either cyclic mechanical stretch or endothelin-1 (ET-1) treatment to induce mechanical and hormonal pro-hypertrophic stress, respectively. P1891A-hiPSC-CMs exhibited a stress-response upon 48 hours of mechanical stretch, that was otherwise absent at 24 hours (Figure S5), resulting in an upregulation of *NPPB* (Figure 7A), whilst levels of *NPPA* remained unchanged (Figure 7B). Expression of *SCN5A* was not influenced by mechanical stretch; however, *SCN5A* was downregulated in the P1891A-hiPSC-CMs in response to the beta blocker metoprolol (Figure 7C). Intriguingly, the combinatorial approach of mechanical stretch and metoprolol significantly increased the expression of the skeletal α-actinin gene, *ACTA1*. In contrast, P1891A-hiPSC-CMs responded similarly to the healthy control hiPSC-CMs following ET-1 treatment, which upregulated *NPPA*, *NPPB*, and *ACTA1* in both hiPSC-CM cell lines, whilst *SCN5A* expression exhibited a trend towards reduction in both lines (Figure 7E-H). This divergent response to different forms of stress suggests that the mechanosensitivity of Na_v_1.5 p.P1891A is altered relative to WT Na_v_1.5.

**Figure 7.**
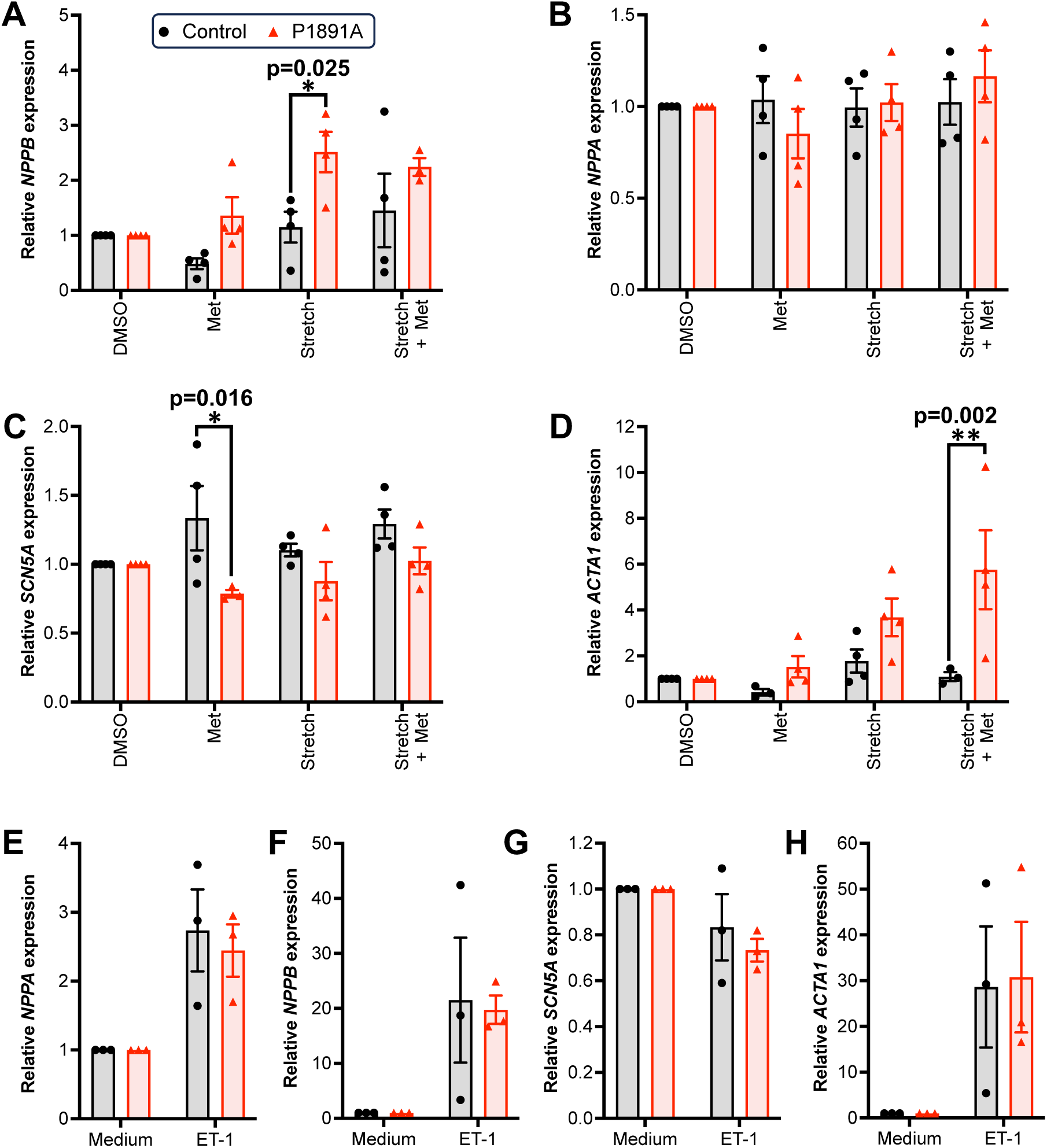
P1891A-hiPSC-CMs exhibit a unique stress-response profile. qPCR analysis of **(A)** *NPPB*, **(B)** *NPPA*, **(C)** *SCN5A*, and **(D)** *ACTA1* gene expression in healthy control-(black bars) and P1891A-hiPSC-CMs (red bars) following 48-hour treatment with either 0.1% DMSO, 10 µM metoprolol (Met), cyclic mechanical stretch, or cyclic mechanical stretch with metoprolol. Data are normalised to the cell-line specific DMSO control and are presented as mean ± SEM, N=4 independent hiPSC-CM differentiations. The data was analysed via a two-way ANOVA followed by a Bonferroni’s multiple comparison test: * P<0.05 and **P<0.01. qPCR analysis of **(E)** *NPPA*, **(F)** *NPPB*, **(G)** *SCN5A*, and **(H)** *ACTA1* gene expression in healthy control-(black bars) and P1891A-hiPSC-CMs (red bars) following either 48-hour culture in RB^+^ medium or with 100 nM endothelin-1 (ET-1) treatment. Data are normalised to the cell-line specific media control and presented as mean ± SEM, N=4 independent hiPSC-CM differentiations.

### Engineered heart tissues derived from P1891A-hiPSC-CMs failed to fully condense and exhibited weaker contractile parameters

Finally, given the phenotypic differences between the two cell lines were modest in 2D cultures, we investigated the effects of the *SCN5A*-P1891A mutation in a 3D fibrin-based contractile cardiac model termed engineered heart tissues (EHTs). In this setting, hiPSC-CMs experience uniaxial static stress when encapsulated within a fibrin hydrogel that is suspended between two silicone posts ^32, 33^. This causes hiPSC-CMs to migrate within the hydrogel in order to align in the direction of the stress, thereby facilitating cell-cell contacts. In turn, the hydrogel becomes thinner, a process referred to as either condensation ^34^ or compaction ^35^.

All EHTs derived from the healthy control hiPSC-CM line underwent full condensation (Figure 8Ai, 8B, and 8C); however, those derived from P1891A-hiPSC-CMs (herein P1891A-EHTs) displayed two distinct phenotypes: partial or full condensation (Figure 8Aii and 8Aiii). Whilst the partially condensed EHTs (67% of the P1891-EHTs, Figure 8B) continued to condense over the recorded 60-day post-fabrication period (Figure 8C), they remained thicker than their age-matched healthy control counterparts. On the other hand, fully condensed P1891-EHTs (33% of the P1891A-EHTs) condensed rapidly within 7 days of fabrication, far quicker than the healthy control EHTs (Figure 8C). By day 30, the P1891A-EHTs had become excessively thin (Figure 8Aiii and 8C), resulting in the fracturing and collapse of the hydrogel in advance of the 60-day end point, thus preventing assessment of contractile properties for the full duration of the experiment.

**Figure 8.**
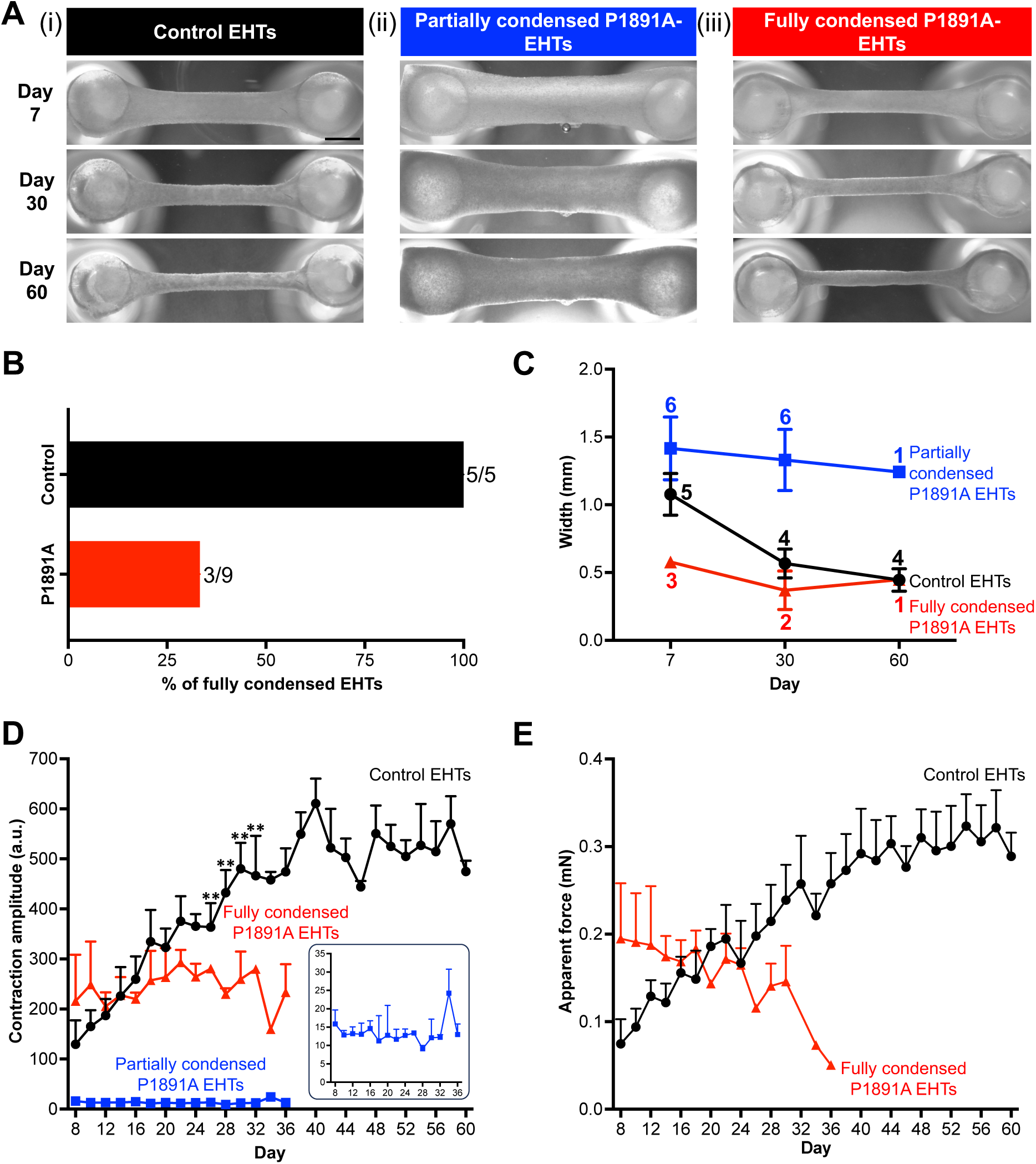
A subset of EHTs derived from P1891A-hiPSC-CMs do not fully condense: Representative light microscopy images of EHTs derived from: (i) healthy control hiPSC-CMs (healthy control EHTs), (ii) P1891A-hiPSC-CMs that underwent partial condensation (partially condensed P1891A-EHTs), and (iii) fully condensed P1891A-EHTs imaged at 7-, 30-, and 60-days post-fabrication. Scale bar represents 1 mm. **(B)** The percentage of EHTs derived from healthy control- or P1891A-hiPSC-CMs that fully condensed. The fraction of fully condensed EHTs is displayed next to each bar. **(C)** Quantification of EHT width 7-, 30-, and 60-days post-fabrication. Healthy control EHTs (black trace), partially condensed P1891A-EHTs (blue trace), fully condensed P1891A-EHTs (red trace). Data are presented as mean ± SEM. N numbers are listed next to each data point and vary due to EHTs fracturing with age. Quantification of the **(D)** contraction amplitude and **(E)** apparent contractile force generated by the different EHTs. Inset: the contraction amplitude of partially condensed P1891A EHTs that were only recorded until day 36 as they did not enhance their contractile parameters. Plotted data are averaged from two consecutive days. Data are presented as mean + SEM. N=3-5 EHTs arising from hiPSC-CMs from unique differentiations. Data was analysed with a mixed-effects model followed by a Tukey’s post-hoc test: ** P<0.01.

Intriguingly, despite forming condensed structures quicker than the healthy control EHTs (Figure 8C), the fully condensed P1891A-EHTs demonstrated significantly weaker contraction amplitudes (Figure 8D) compared to the healthy control EHTs. Indeed, whilst the control EHTs demonstrated a positive correlation between post-fabrication age and contraction amplitude, the P1891A-EHT underwent age-associated functional decline (Figure 8D). This observation was similarly mirrored by the apparent force generated by the healthy control EHTs (Figure 8E). Although the P1891A-EHTs displayed a trend towards greater initial apparent force, in line with their capacity to condense quicker than other EHTs (Figure 8C), this potential reduced over time suggesting an inability of the encapsulated P1891A-hiPSC-CMs to sustain or enhance contractility in this 3D model. In keeping with their failure to fully condense the hydrogel, the partially condensed P1891A-EHTs did not elicit robust contraction or apparent force and instead exhibited uncoordinated, low-magnitude twitching (Figure 8D - 8E, Supplementary Videos 1 - 3)

## Discussion

The potential role of Na_v_1.5 mutations in LVHT pathophysiology has thus far largely focused upon its involvement in the context of arrhythmia induction ^18, 36^. However, LVHT is a highly complex condition with pathophysiological features extending beyond electrophysiological parameters. Utilising hiPSC technology, electrophysiological assessment, advanced proteomics, in vitro stimulation, and 3D cardiac tissues, we extensively characterised a single novel missense mutation (*SCN5A*-P1891A) from an LVHT patient without concomitant disease, thereby providing novel insights into the role of the mutated Na_v_1.5 sodium channel in LVHT aetiology.

Our electrophysiological assessment of the P1891A-hiPSC-CMs revealed reduced Na^+^ current densities, increased Na^+^ window current, and arrhythmogenicity; however, AP parameters were not perturbed. Although the P1891A mutation did not fully shift Na_v_1.5 inactivation toward depolarised potentials, the incomplete inactivation started at approximately -70 mV resulting in a larger window current. The probability of Na_v_1.5 opening during the end phase of cardiac repolarisation is determined by the crossover of the activation and inactivation curves at which point, a fraction of Na_v_1.5 channels have recovered from inactivation and may reopen ^37^. A larger window current may potentially increase the intracellular Na^+^ concentration by generating larger Na^+^ influx that offsets the balance between depolarising and repolarising currents in favour of depolarisation. In turn, this may elicit arrhythmias ^38^ and could explain the enhanced occurrence of arrhythmias in the P1891A-hiPSC-CMs.

Other C-terminal mutations of Na_v_1.5 attribute their arrhythmic phenotype to perturbed inactivation parameters. To this end, the Na_v_1.5 p.V1951L GOF mutation, identified in an LVHT patient presenting with concomitant NSVT, reduces the slow inactivation of the channel ^36^, a hallmark known to be associated with enhanced arrhythmias ^39^. Similarly, the Na_v_1.5 p.H1849R GOF mutation, implicated in LQT3, prevents FGF13 from binding to Na_v_1.5, resulting in a slower rate of channel inactivation, leading to a prolonged APD that manifests as arrhythmogenic afterdepolarisations ^40^. Intriguingly, this is also associated with an increase in the Na^+^ current density. However, the APD of the P1891A-hiPSC-CMs was unperturbed, suggesting this was not responsible for the enhanced arrhythmogenicity. FGF12 also binds to the C-terminal of Na_v_1.5, where it reduces the voltage dependence of inactivation ^27^. The Na_v_1.5 p.D1790 mutation, also associated with LQT3, prevents Na_v_1.5 from interacting with FGF12 ^27^. FGF12 and FGF13 bind to the C-terminal of Na_v_1.5 at residues 1773-1832 and 1849^27, 28^, respectively. Our proteomics analysis revealed that the p.P1891A mutation abrogated interactions with both FGFs despite being downstream of their binding sites. This is likely because the amino acid immediately upstream of the P1891A mutation, E1890, plays an essential role in binding to FGF13 by establishing a salt bridge with lysine (K)14 of FGF13 ^41^. The P1891A mutation may therefore alter the tertiary structure of this binding pocket, thereby preventing this known interaction.

Notably, the Na^+^ current density can be modulated irrespective of the gating properties of Na_v_1.5. Indeed, its PPIs, including those with sarcomeric Z-disk proteins such as α-actinin-2, upregulate the Na^+^ current density ^23^, whilst the inability to interact with ankyrin-G, responsible for localising the Na^+^ channel to the cardiomyocyte T-tubules, manifests as Brugada syndrome ^42^. Thus, in addition to understanding the interactions that were perturbed via the P1891A mutation, we also conducted advanced proteomics on the wild type Na_v_1.5 protein to ascertain novel interactors at both the N and C terminus. Of these, several were implicated in mitochondrial processes. This follows the recent discovery of the Na_v_1.5-mitochondrial couplon, whereby the interaction of Na_v_1.5 with the Z-disks ensures proximity to subsarcolemmal mitochondria, enabling Na^+^ influx and function of the Na^+-^Ca^2+^ exchanger (NCX) to prevent calcium-mediated mitochondrial injury ^24^. This further highlights the importance of the Na_v_1.5 beyond its role in electrogenesis.

To this end, cardiac development in zebrafish depends on the *SCN5A* orthologue, *scn5Lab*, which drives cardiomyocyte proliferation prior to detection of the Na^+^ current ^25^. In the mouse embryo, LVHT-causing mutations have demonstrated disparate effects on cardiomyocyte proliferation, either downregulating proliferation within the compact layer at E12.5 ^6^ or upregulating proliferation of the trabecular layer at E15.5 ^7^. This results in a thin compact layer or a thick trabecular, respectively, both of which perturb the ratio of the two layers and results in LVHT. These observations challenge the original non-compaction hypothesis, curated in chick embryos, that centres around a thick trabecular layer emerging from a failure of the developing ventricular trabeculae to compact ^43^. To the best of our knowledge, our data is the first to associate Na_v_1.5 mutations with enhanced proliferation of human cardiomyocytes. Although LVHT largely arises due to genetic factors in utero ^6, 7^, environmental stimuli such as excessive preload, as can be experienced during pregnancy, can cause a transient increase in LV trabeculation in patients who carry a LVHT-causing mutation ^44^. This process would need to be driven by the proliferation of cardiomyocytes within the trabecular layer that would in turn shift the trabecular-to-compact ratio and manifest as adult-onset LVHT.

With this in mind, Na_v_1.5 itself is mechanosensitive ^31^ and thus responsive to preload. This ability of the Na^+^ channel to respond to mechanical stimulation, a process that appears to be mediated by the channel pore ^45^, is critical to its function beyond electrogenesis. Indeed, the Na_v_1.5 p.G615E mutation attenuates mechanosensitivity leading to LQT3 independent of the voltage-dependent function of the channel ^46^. Our proteomics analysis identified the Golgi-residing ion channel, TMEM87A as a novel interactor of Na_v_1.5. Whilst this channel has also been reported to be mechanosensitive ^47^ more recent observations have suggested it modulates the function of other mechanosensitive ion channels ^48^. Therefore, our proteomics data potentially reveals a new regulator of WT Na_v_1.5 that is not perturbed by the P1891A mutation. However, despite this interaction, the P1891A-hiPSC-CMs displayed an altered stress response following cyclic mechanical stretch, whereas no such response was observed with ET-1-mediated stress. Indeed, stretch or stretch with metoprolol administration caused a pronounced upregulation of *NPPB* and *ACTA1* in P1891A-hiPSC-CMs, an expression pattern observed in hypertrophy and heart failure, respectively, During heart failure, adult cardiomyocytes dedifferentiate and adopt a foetal-like gene expression programme ^51^. During early cardiomyocyte differentiation, hiPSC-CMs are also foetal-like and preferentially express the non-cardiac isoforms of the Na^+^ channel, particularly the neuronal *SCN2A*, with *SCN5A* not becoming the dominant isoform until differentiation day 20 ^52^. As in early development, *SCN5A* expression is also lower during ageing ^53^. Administration of metoprolol alone to the P1891A-hiPSC-CMs resulted in a trend towards reduced *SCN5A* expression. Taken together, P1891A-hiPSC-CMs transition towards a heart failure-like state following cyclic mechanical stimulation.

This perturbed response to cyclic mechanical stretch may explain the divergent phenotype observed with the P1891A-EHTs. Indeed, it is plausible the P1891A-hiPSC-CMs similarly fail to respond to static stress, thereby overwhelmingly preventing condensation of the fibrin hydrogel. Although a small subset of P1891A-EHTs rapidly condensed and became contractile, they demonstrated an age-associated functional decline, consistent with the heart failure-like gene expression profile adopted by the P1891A-hiPSC-CMs following cyclic mechanical stretch. Moreover, the effects of the P1891A mutation may be further exacerbated in this 3D model. To this end, EHTs derived from healthy hiPSC-CMs generate a greater Na^+^ current than their 2D monolayer controls ^52^. It is therefore possible that the Na^+^ current density of the P1891A-hiPSC-CMs is even further reduced in the context of the EHT.

Direct comparison with other relevant mutation-carrying EHTs has proven challenging given a number of recent studies characterising *SCN5A* mutations ^54^, or those associated with LQT syndrome ^55^, or Brugada syndrome ^56^ have been limited to 2D monolayers and have not yet utilised the EHT platform. *SCN5A* mutations have, however, been evaluated in heart-on-a-chip biowires that similarly demonstrate a reduction in the force generation capacity of *SCN5A* mutation-carrying hiPSC-CMs ^57^.

Taken together, our in-depth, multi-parametric characterisation of the novel *SCN5A*-P1891A missense mutation reveals a potential causative effect of *SCN5A* mutations in LVHT aetiology. Our study demonstrated that the P1891A-hiPSC-CMs mirrored the arrhythmogenicity of the mutation-carrying donor, possibly due to a larger Na^+^ window current. Beyond electrophysiological considerations, the P1891A mutation perturbed hiPSC-CM function, affecting the response of P1891A-hiPSC-CMs to both mitogenic stimuli and cyclic mechanical stimulation, as well as static mechanical stimulation in the 3D EHT. In turn, this heavily disrupted 3D model provides an excellent tool for further in-depth mechanistic research as well as drug development studies for the treatment of mutated Na_v_1.5-mediated LVHT.

## Non-standard Abbreviations and Acronyms

LVHT: Left ventricular hypertrabeculation
LVNC: Left ventricular noncompaction
hiPSC-CMs: Human induced pluripotent stem cell-derived cardiomyocytes
MAC: Multiple approaches combined-tagged
PPIs: Protein-protein interactions
HCIs: High-confidence interactions
EHT: Engineered heart tissue

## Acknowledgements, Sources of Funding, & Disclosures

## a) Acknowledgments

We thank Annika Korvenpää, Henna Lappi, and Markus Haponen for technical support; Faiza Moureen for calcium imaging; Nuutti Lahtinen for cell line authentication; and Saana Pohjavaara for preparing DNA samples for Sanger Sequencing. The authors also acknowledge the Biocentre Finland-Finnish Electrophysiology Platform, Tampere Imaging Facility - light microscopy core, and Tampere University’s iPSC core facility for their services.

## b) Sources of Funding

QAM: The Finnish Foundation for Cardiovascular Research, The Finnish Cultural Foundation LP: The Finnish Foundation for Cardiovascular Research, The Finnish Cultural Foundation KAS: Research Council of Finland (Centre of Excellence in Body-on-Chip Research), Finnish Foundation for Cardiovascular Research, Sigrid Jusélius Foundation.

VT: Research Council of Finland (projects 321564; 328909; 353109), The Finnish Foundation for Cardiovascular Research, Sigrid Jusélius Foundation, Päivikki and Sakari Sohlberg Foundation.

## c) Disclosures

Subsets of the data were presented at the European Society of Cardiology (ESC) Congress 2024 and the 48th ESC Working Group on Cardiac Cellular Electrophysiology (EWGCCE) Meeting 2024.

## Supplemental Material

Expanded Materials and Methods

Figures S1–S5

Tables S1–S3

Videos S1–S3

Major Resources Table

References 20, 22, 26, 32, 58–67

## Novelty and Significance

### What is known?

- Left ventricular hypertrabeculation (LVHT) is a poorly understood cardiac disease arising from genetic or environmental factors.
- Mutations in *SCN5A*, the gene encoding the cardiac sodium channel Na_v_1.5, have been linked to several cardiac diseases but no causal relationship with LVHT has been established.
- The functions of Na_v_1.5 are modified by protein-protein interactions through which the channel may participate in cellular processes other than the regulation of the action potential.

### What new information does this article contribute?

- We report that a novel *SCN5A*-P1891A mutation is associated with LVHT in a Finnish family and alters Na_v_1.5 electrophysiology in human induced pluripotent stem cell-derived cardiomyocytes carrying the mutation (P1891A-hiPSC-CMs).
- We show that the P1891A-hiPSC-CMs exhibited an altered response to mitogenic and mechanical stimuli, largely impairing the formation and function of 3D engineered cardiac tissues.
- We report the complete protein interactome of both wild type and mutated Na_v_1.5 and reveal disrupted interactions with FGF12 and FGF13 as a probable cause for the disease phenotype.

LVHT is a complex cardiac disease with both genetic and environmental contributors, although the underlying mechanisms are not yet fully understood. Whilst *SCN5A* mutations are known to cause ion channel disorders and arrhythmias, their involvement in LVHT aetiology has not yet been established. Here, we report a novel *SCN5A*-P1891A mutation associated with LVHT in a Finnish family. We provide the first comprehensive analysis of the complete protein interactome of Na_v_1.5 and established how this was perturbed by the P1891A mutation. This insight holds profound potential in enhancing our understanding of this sodium channel, particularly its non-electrophysiological involvement. Indeed, we demonstrated, for the first time, that an *SCN5A* mutation enhances human cardiomyocyte proliferation. Further, an altered stress response following cyclic mechanical stretch indicated dysregulated channel mechanosensitivity. This likely explained why P1891A-hiPSC-CMs failed to respond to static stress in 3D cardiac tissues and were unable to generate functional tissues with robust contractility. This work not only advances our understanding of LVHT aetiology but, through the novel insight into the sodium channel itself, reveals how its mutations can affect different cardiomyocyte processes. The establishment of a robust 3D functional model of LVHT allows for future studies that explore therapeutic interventions that address both the electrical and mechanical aspects of the disease.

